# Chromosome-scale genome assembly and *de novo* annotation of *Alopecurus aequalis*

**DOI:** 10.1101/2024.10.09.616956

**Authors:** Jonathan Wright, Kendall Baker, Tom Barker, Leah Catchpole, Alex Durrant, Fiona Fraser, Karim Gharbi, Christian Harrison, Suzanne Henderson, Naomi Irish, Gemy Kaithakottil, Ilia J. Leitch, Jun Li, Sacha Lucchini, Paul Neve, Robyn Powell, Hannah Rees, David Swarbreck, Chris Watkins, Jonathan Wood, Seanna McTaggart, Anthony Hall, Dana MacGregor

## Abstract

*Alopecurus aequalis* is a winter annual or short-lived perennial bunchgrass which has in recent years emerged as the dominant agricultural weed of barley and wheat in certain regions of China and Japan, causing significant yield losses. Its robust tillering capacity and high fecundity, combined with the development of both target and non-target-site resistance to herbicides means it is a formidable challenge to food security. Here we report on a chromosome-scale assembly of *A. aequalis* with a genome size of 2.83 Gb. The genome contained 33,758 high-confidence protein-coding genes with functional annotation. Comparative genomics revealed that the genome structure of *A. aequalis* is more similar to *Hordeum vulgare* rather than the more closely related *Alopecurus myosuroides* and has undergone an expansion of cytochrome P450 genes, a gene family involved in non-target-site herbicide resistance.

## Background and Summary

*Alopecurus aequalis,* commonly known as shortawn foxtail or orange foxtail, is a winter annual or short-lived perennial bunchgrass of the Poaceae family. It is native to at least 55 different countries across the Northern Hemisphere and northern Southern Hemisphere and has been introduced into Australasian regions^1^. *A. aequalis* has emerged as the dominant agricultural weed of winter canola, barley, and wheat only in certain regions of China and Japan despite its widespread distribution^2^. *A. aequalis* can cause significant yield losses; densities of up to 1,560 plants per m^2^ reduce wheat yields by up to 50%^3^. The biology of *A. aequalis,* particularly its robust tillering capacity and high fecundity (with a single plant able to produce over 7,300 highly dispersible seeds), makes it a challenging weed to control^3^. Moreover, the evolution of both target-site resistance (TSR^4,5^) and non-target-site resistance (NTSR^5–7)^ means many of the available chemical control methods are ineffective. Therefore, *A. aequalis* is a formidable challenge to food security and novel and innovative control methods are urgently required.

Another Poaceae grass, *Alopecurus myosuroides* (black-grass), has evolved to occupy similar agroecosystem niches. Like *A. aequalis*, *A. myosuroides* is a weed of winter cereals in China and Japan^8,9^ and surveys have recorded it as present across the Northern Hemisphere^1^. However, *A. myosuroides* has become the predominant agricultural weed in Western European winter wheat and barley, leading to considerable yield losses and economic consequences^10^. These two species have similar but distinct morphologies and growth habits (Figure 1). Like *A. aequalis*, black-grass exhibits widespread multiple-herbicide resistance^10–12^ and the characterized resistance mechanisms are similar between the two species. Both have TSR mutations that alter equivalent amino acids of homologous herbicide target genes^4^ and NTSR correlated with increased xenobiotic-metabolizing enzymes such as cytochrome P450 mono-oxygenases and glutathione s-transferases^6,7^. In black-grass, NTSR is highly heritable with no evidence that it results in a fitness penalty^13^, and it is correlated with increased tolerance to drought and waterlogging stresses^14,15^. There is evidence that some TSR mutations are associated with fitness costs^16^. These characteristics, combined with an ability to compete with crops for essential resources like nutrients, water, and light, mean that when either foxtail species are present in agricultural fields, they significantly reduce crop yields and overall productivity within agroecosystems^3,10,11,14,15^.

**Figure 1:**
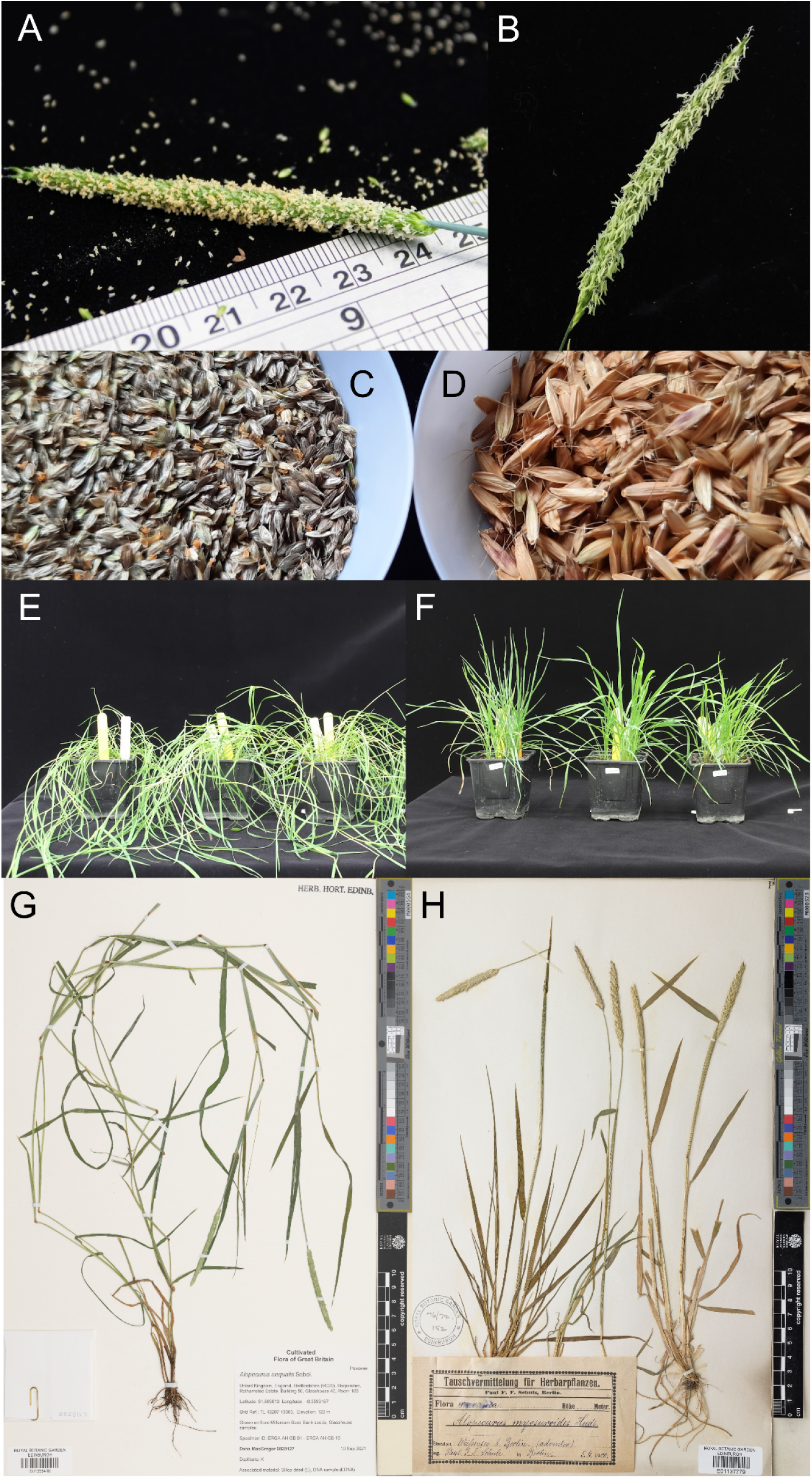
Images highlighting differences in *Alopecurus aequalis* (shortawn foxtail or orange foxtail; A, C, E, & G) and *Alopecurus myosuroides* (black-grass; B, D, F, & H) morphologies. Images show flowering heads (A-B), seeds (C-D), at vegetative growth stage (E-F) and Kew Herbarium images (G-H). The flower spike of *A. aequalis* (Figure 1A) has a blunt end rather than the tapered end in *A. myosuroides* (B). While both have single flowered spikelets, the anthers of *A. aequalis* are shorter compared to *A. myosuroides* (Figure 1B). The mature seeds of these two species are easily distinguished (Figure 1C *A. aequalis*, Figure 1D *A. myosuroides*). *A. aequalis* has a more prostrate growth (Figure 1E) while *A. myosuroides* is more upright (Figure 1F). These differences can be seen in Herbarium images from Royal Botanic Garden Edinburgh (https://data.rbge.org.uk/search/herbarium/) for *A. aequalis* (Barcode: E01358418, Figure 1G), and *A. myosuroides* (Barcode: E01137779, Figure 1H).

Despite geographic isolation and 7.4 million years of divergence^17^, these two species have evolved similar herbicide resistance mechanisms and have become problematic in similar winter crops. It is not yet understood whether similarities between these two species are the result of parallel evolution. This lack of direct comparison is in part due to lack of genomic data for either species. Recently, two reference genomes have been produced for biotypes of *A. myosuroides* that are sensitive to all tested herbicides^18,19^. We therefore set out to generate a genome of similar quality for *A. aequalis* as part of the European Reference Genome Atlas^20^ (ERGA) pilot programme, which aims to empower research communities to expand the taxonomic coverage of genomic resources to address continent-scale questions at the genomic level.

Here we report a *de novo* annotated, chromosome-level assembly of *A. aequalis.* PacBio HiFi reads (32.9x coverage) were used to assemble the genome resulting in a contig assembly of 2.9 Gb with a contig-N50 of 374.7 Mb. The assembled size was identical to the estimated genome size from *k-mer* based methods. Omni-C reads (56.7x coverage) were used to anchor and orient the contigs into seven pseudomolecules. Protein-coding genes were annotated using REAT^21^ an evidence-guided pipeline, making use of RNA-Seq alignments, transcript assemblies from Iso-Seq reads and alignment of protein sequences. In total, 33,758 high-confidence protein-coding genes were identified. Whole genome alignment between *A. aequalis*, *A. myosuroides* and *Hordeum vulgare* indicated that the genome structure of *A. aequalis* was more similar to *H. vulgare* than to the more closely related *A. myosuroides.* The number of cytochrome P450 genes, previously identified as being important in the evolution of non-target-site herbicide resistance^22^, was of similar magnitude to that found in *A. myosuroides* indicating that this important weed has also undergone an expansion in cytochrome P450 genes. This genomic resource provides a much-needed foundation for investigating the molecular mechanisms underlying weedy traits, such as widespread multiple-herbicide resistance, in *Alopecurus aequalis*, and will be an invaluable tool for the research community in devising more effective weed management strategies.

## Methods

### *Alopecurus aequalis* plants and materials

Seeds of *Alopecurus aequalis* (orange foxtail), donated by a private Individual in 2014 to the Royal Botanic Gardens, Kew Millennium Seed Bank (Serial Number 828127), were used for genome sequencing and annotation. While the collection location was not recorded, *A. aequalis* is not an agricultural weed in the UK, making it unlikely that it was collected from an active agricultural field. Consequently, these seeds were considered a herbicide-naive *A. aequalis* genotype. To establish a seed stock, 26 plants were grown and allowed to intrabreed in isolation for one generation at Rothamsted Research. For genome size estimation by flow cytometry, fresh leaf material from four individual plants (ID 828127A, B, C and D) of this second generation were analysed using the fluorochrome propidium iodide with the ‘one-step method’^23^. Nuclei were isolated in the LB01 nuclei isolation buffer^24^ and *Petroselinum crispum* ‘Champion Moss Curled’ was used as the internal calibration standard with an assumed 2C-value of 4.5 pg^25^. The mean relative fluorescence of nuclei from *A. aequalis* and *P. crispum* were used to estimate the genome size of *A. aequalis* using the following equation: 2C-value of *A. aequalis* (pg) = (Mean peak position of *A. aqualis* / mean peak position of *P. crispum*) ✕ 4.5.

To generate sufficient material for genome sequencing, a single plant (ID 828217A) was grown and vegetatively cloned twice following protocols described in the supplementary material in Cai et al. (2023)^18^. DNA for genome sequencing was isolated using the protocols described below from young leaves and meristem material from plants that had been dark-adapted for five days, flash frozen in liquid nitrogen, and stored at -80°C until shipping on dry ice. Similarly, RNA for Iso-Seq and RNA sequencing for annotation was sourced from flag leaves and flowering heads from clones of plants 828217A and 828217B. Additional RNA came from bulked shoot or root material derived from five technical replicates of 5-8 plants from the 828217 seed stock. These plants were grown under sterile hydroponic conditions, as per supplementary material of Cai et al. (2023), for a total of 42 days before separating root or shoot material from the seed. The flash frozen material was stored at -80°C until it was shipped on dry ice for processing. One clone from 828217A was allowed to flower and preserved by preparing a herbarium voucher which is stored at the Royal Botanic Garden Edinburgh (E) (Figure 1G, https://data.rbge.org.uk/herb/E01358418)

### DNA extraction

HMW DNA extraction was performed using the Nucleon PhytoPure kit, with a slightly modified version of the manufacturer’s protocol. One gram of snap-frozen leaf material was ground under liquid nitrogen for 10 minutes. The tissue powder was thoroughly resuspended in Reagent 1 using a 10mm bacterial spreader loop until the mixture appeared completely homogeneous, at which point 4µl of 100mg/ml RNase A (Qiagen cat no. 19101) was added and mixed again.

After incubation on ice, 200µl of resin was added along with chloroform. The chloroform extraction was followed by a phenol:chloroform extraction. Here, an equal volume of 25:24:1 phenol:chloroform:isoamyl alcohol was added to the previous upper phase, mixed gently at 4°C for 10 minutes and then centrifuged at 3000g for 10 minutes. The upper phase from this procedure was then transferred to another 15ml Falcon tube and precipitation proceeded as recommended by the manufacturer’s protocol. The final elution was left open in a chemical safety cabinet for two hours to allow residual phenol and ethanol to evaporate, and the DNA sample was left at room temperature overnight before further processing.

### PacBio HiFi library preparation and sequencing

In total, 65µg of genomic DNA was split into 5 aliquots and manually sheared with the Megaruptor 3 instrument (Diagenode, P/N B06010003), according to the Megaruptor 3 operations manual, loading each aliquot of 150µl at 21ng/µl, with a shear speed of 31. Each sheared aliquot underwent AMPure® PB bead (PacBio®, P/N 100-265-900) purification and concentration before undergoing library preparation using the SMRTbell® Express Template Prep Kit 2.0 (PacBio®, P/N 100-983-900). The HiFi libraries were prepared according to the HiFi protocol version 03 (PacBio®, P/N 101-853-100) and the final libraries were size fractionated using the SageELF® system (Sage Science®, P/N ELF0001), 0.75% cassette (Sage Science®, P/N ELD7510), running on a 4.55hr program with 30µl of elution buffer per well post elution. The libraries were quantified by fluorescence (Invitrogen Qubit™ 3.0, P/N Q33216) and the size of the library fractions were estimated from a smear analysis performed on the FEMTO Pulse® System (Agilent, P/N M5330AA).

The loading calculations for sequencing were completed using the PacBio® SMRT®Link Binding Calculator 10.2. Sequencing primer v2 was annealed to the adapter sequence of the HiFi libraries. The libraries were bound to the sequencing polymerase with the Sequel® II Binding Kit v2.0. Calculations for primer and polymerase binding ratios were kept at default values for the library type. Sequel® II DNA internal control 1.0 was spiked into each library at the standard concentration prior to sequencing. The sequencing chemistry used was Sequel® II Sequencing Plate 2.0 (PacBio®, P/N 101-820-200) and the Instrument Control Software v 10.1. The libraries were sequenced on the Sequel IIe across 15 Sequel II SMRT®cells 8M. The parameters for sequencing per SMRT cell were diffusion loading, 30-hour movie, 2-hour immobilisation time, 4-hour pre-extension time, 60-80pM on plate loading concentration. We generated 11.7 million PacBio HiFi reads (227 Gb of sequence), corresponding to a haploid genome coverage of 32.9x. The average HiFi read length was 19.4 Kbp.

### Dovetail Omni-C library preparation and sequencing

Sample material for the Omni-C library preparation was 300mg of the youngest leaf and meristem material from plants dark adapted for 5 days then flash frozen at harvest. The Omni-C library was prepared using the Dovetail® Omni-C® Kit (SKU: 21005) according to the manufacturer’s protocol^26^.

Briefly, the chromatin was fixed with disuccinimidyl glutarate (DSG) and formaldehyde in the nucleus. The cross-linked chromatin sample was then digested *in situ* with 0.05µl DNAse I. Following digestion, the cells were lysed with SDS to extract the chromatin fragments and the chromatin fragments were bound to Chromatin Capture Beads. Next, the chromatin ends were repaired and ligated to a biotinylated bridge adapter followed by proximity ligation of adapter-containing ends. After proximity ligation, the crosslinks were reversed, the associated proteins were degraded, and the DNA was purified before conversion into a sequencing library (NEBNext® Ultra™ II DNA Library Prep Kit for Illumina® (E7645)) using Illumina-compatible adaptors (NEBNext® Multiplex Oligos for Illumina® (Index Primers Set 1) (E7335)).

Biotin-containing fragments were isolated using streptavidin beads prior to PCR amplification. The Omni-C library was quantified by qPCR using a Kapa Library Quantification Kit (Roche Diagnostics 7960204001). The pool was diluted to 2nM and denatured using 2N NaOH before diluting to 20pM with Illumina HT1 buffer. The denatured pool was loaded on an Illumina MiSeq Sequencer for quality control with a 300 cycle MiSeq Reagent Kit v2 (Illumina MS-102-2002) at 10pM concentration with a 1% phiX control v3 spike (Illumina FC-110-3001). The MiSeq was run using control software version 4.0, RTA v1.18.54.430, and the data was demultiplexed and converted to fastq using bcl2fastq2. Following quality control analysis, the library was sequenced on one lane of a 300 cycle NovaSeq X Series 10B Reagent Kit (Illumina 20085594). For this run, the library was diluted down to 0.75nM using EB (10mM Tris pH8.0) in a volume of 40µl before spiking in 1% Illumina phiX Control v3. This was denatured by adding 10µl 0.2N NaOH and incubating at room temperature for 5 mins, after which it was neutralised by adding 150µl of Illumina’s preload buffer, of which 160µl was loaded onto the NovaSeq X Plus for sequencing. The NovaSeq X Plus was run using control software version 1.0.1.7385 and was set up to sequence 150bp paired-end reads. The data was demultiplexed and converted to fastq using bcl2fastq2. We generated 1.28 billion reads representing 56.7x haploid genome coverage.

### Illumina RNA-Seq library preparation and sequencing

Five root samples and five leaf samples were used to generate RNA-Seq libraries. This was performed on the Perkin Elmer (formerly Caliper LS) Sciclone G3 (PerkinElmer PN: CLS145321) using the NEBNext Ultra II RNA Library prep for Illumina kit (NEB#E7760L) NEBNext Poly(A) mRNA Magnetic Isolation Module (NEB#E7490L) and NEBNext Multiplex Oligos for Illumina® (96 Unique Dual Index Primer Pairs) (E6440S/L) at a concentration of 10µM. One µg of RNA was purified to extract mRNA with a Poly(A) mRNA Magnetic Isolation Module. Isolated mRNA was then fragmented for 12 minutes at 94°C, and converted to cDNA. NEBNext Adaptors were ligated to end-repaired, dA-tailed DNA. The ligated products were subjected to a bead-based purification using Beckman Coulter AMPure XP beads (A63882) to remove unligated adaptors. Adaptor Ligated DNA was then enriched by 10 cycles of PCR (30 secs at 98°C, 10 cycles of: 10 secs at 98°C _75 secs at 65°C _5 mins at 65°C, final hold at 4°C). The size of the resulting libraries was determined using a Perkin Elmer DNA High Sensitivity Reagent Kit (CLS760672) with DNA 1K / 12K / HiSensitivity Assay LabChip (760517) and the concentration measured with a Quant-iT™ dsDNA Assay Kit, high sensitivity (Plate Reader) assay from ThermoFisher (Q-33120). The final libraries were pooled equimolarly and quantified by qPCR using a Kapa Library Quantification Kit (Roche Diagnostics 7960204001).

The pool was diluted down to 0.5nM using EB (10mM Tris pH8.0) in a volume of 18µl before spiking in 1% Illumina phiX Control v3. This was denatured by adding 4µl 0.2N NaOH and incubating at room temperature for 8 mins, after which it was neutralised by adding 5µl 400mM tris pH 8.0. A master mix of DPX1, DPX2, and DPX3 from Illumina’s Xp 2-lane kit was made and 63µl added to the denatured pool leaving 90µl at a concentration of 100pM. This was loaded onto a single lane of the NovaSeq SP flow cell using the NovaSeq Xp Flow Cell Dock before loading onto the NovaSeq 6000. The NovaSeq was run using NVCS v1.7.5 and RTA v3.4.4 and was set up to sequence 150bp PE reads. The data was demultiplexed and converted to fastq using bcl2fastq2. A total of 253 million and 234 million reads were generated for root and shoot samples respectively.

### PacBio Iso-Seq library preparation and sequencing

Iso-Seq libraries were generated for root and shoot samples. The five shoot extractions were pooled for the shoot library and a single root sample was used for root. The libraries were constructed starting from 356-482ng of total RNA per sample. Reverse transcription cDNA synthesis was performed using NEBNext® Single Cell/Low Input cDNA Synthesis & Amplification Module (NEB, E6421). Each cDNA sample was amplified with barcoded primers for a total of 12 cycles. Each library was prepared according to the guidelines laid out in the Iso-Seq protocol version 02 (PacBio, 101-763-800) using SMRTbell express template prep kit 2.0 (PacBio, 102-088-900). The library pool was quantified using a Qubit Fluorometer 3.0 (Invitrogen) and sized using the Bioanalyzer HS DNA chip (Agilent Technologies, Inc.).

The loading calculations for each Iso-Seq library were calculated using the PacBio SMRTlink Binding Calculator v.10.2.0.133424 and prepared for sequencing according to the library type. Sequencing primer v4 was annealed to the Iso-Seq library pool and complexed to the sequencing polymerase with the Sequel II binding kit v2.1 (PacBio, 101-843-000). Calculations for primer to template and polymerase to template binding ratios were kept at default values for the library type. Sequencing internal control complex 1.0 (PacBio, 101-717-600) was spiked into the final complex preparation at a standard concentration before sequencing for all preparations. The sequencing chemistry used was Sequel® II Sequencing Plate 2.0 (PacBio®, 101-820-200) and the Instrument Control Software v10.1.0.125432.

Each Iso-Seq library was sequenced on the Sequel IIe instrument with one Sequel II SMRT®cell 8M cell per library. The parameters for sequencing per SMRTcell were diffusion loading, 30-hour movie, 2-hour immobilisation time, 2-hour pre-extension time, 60pM on plate loading concentration.

We generated 3.9 million CSS reads for the root sample and 3.8 million CCS reads for the shoot sample. These were filtered to identify Full-Length Non-Concatamer (FLNC) reads resulting in 3.6 and 3.4 million FLNC reads for root and shoot, respectively. Clustering these individually generated 258,543 transcripts for root and 212,332 transcripts for shoot. In addition, root and shoot FLNC reads were combined and clustered to generate a set of 410,286 transcripts.

### Genome survey

A previous study reported that *A. aequalis* has seven chromosomes^27^. Prior to genome sequencing, we first estimated genome size using flow cytometry for four individual plants of *A. aequalis* line 828217 which gave a mean estimated genome size of 3.45 Gb/1C (Table 1).

**Table 1:**
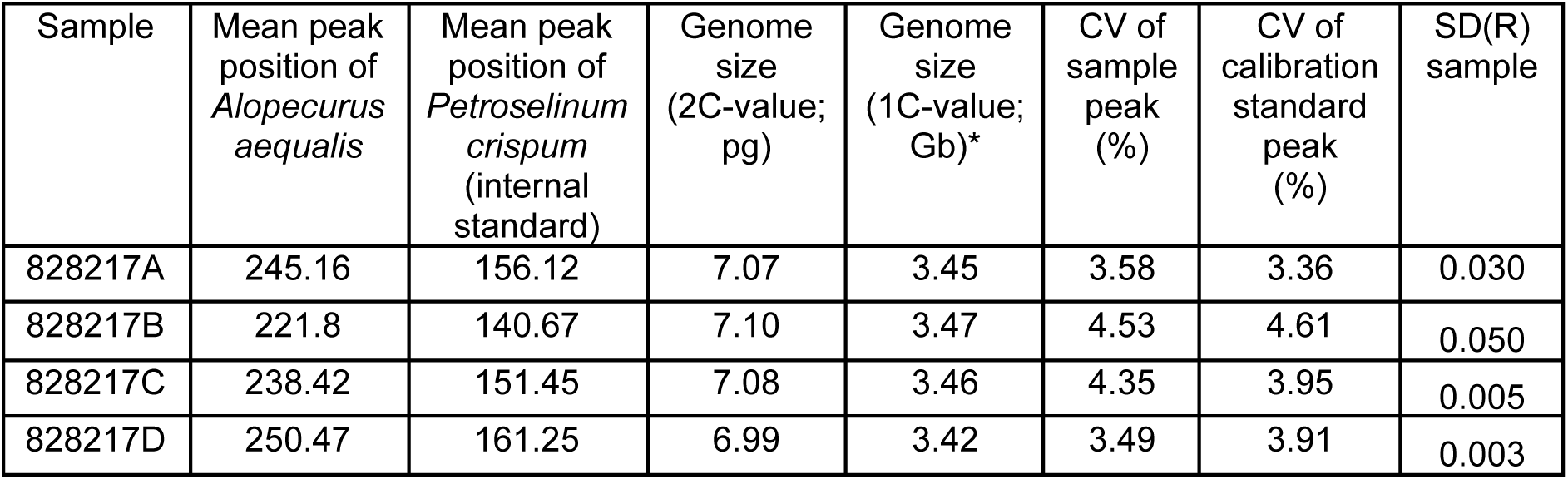
Genome size estimation by flow cytometry for four individual plants of *Alopecurus aequalis* line 828127. (*Flow cytometry estimates in pg were converted to Gb using the conversion factor 1 pg = 0.978 Gb^28^). “CV” is the coefficient of variation for the peaks in our flow cytometry data which are an indication of the reliability of the genome size estimate and “SD(R)” is the standard deviation of the genome size estimate.

We used FastK^29^ (v1.1) to count k-mers (k=31) in the HiFi reads, then GenomeScope.FK (based on GenomeScope 2.0^30^) to explore genome characteristics. In order to estimate genome size, the haploid k-mer coverage was set to 33.

`FastK -k31 -T16 -M200 -P. -Ntmp_fastk reads.fastq.gz`

`Histex -G reads.hist > hist_all.out`

`GeneScopeFK.R -i hist_all.out -o all_out -k 31 --kmercov 33`

The estimated genome size from GenomeScope was 2.9 Gb (Figure 2), which is 0.55 Gb less than the flow cytometry estimate of 3.45 Gb . However, genome size estimates using k-mers are often smaller than those obtained by flow cytometry due to the challenges of assembling the repetitive fraction of the genome. Computational methods try to avoid overestimating genome size by ignoring high frequency k-mers which come from a mix of real repeats and artifacts such as organelles or small contaminants. GenomeScope estimated the heterozygous percentage of the genome as 0.058% and the k-mer profile shows a large single peak centered around coverage 73 representing the homozygous part of the genome. There is a slight deviation from the model on the left side of the peak (marked with a red arrow) representing the heterozygous part of the genome. This indicates the genome is less heterozygous than expected for an outcrossing species.

**Figure 2:**
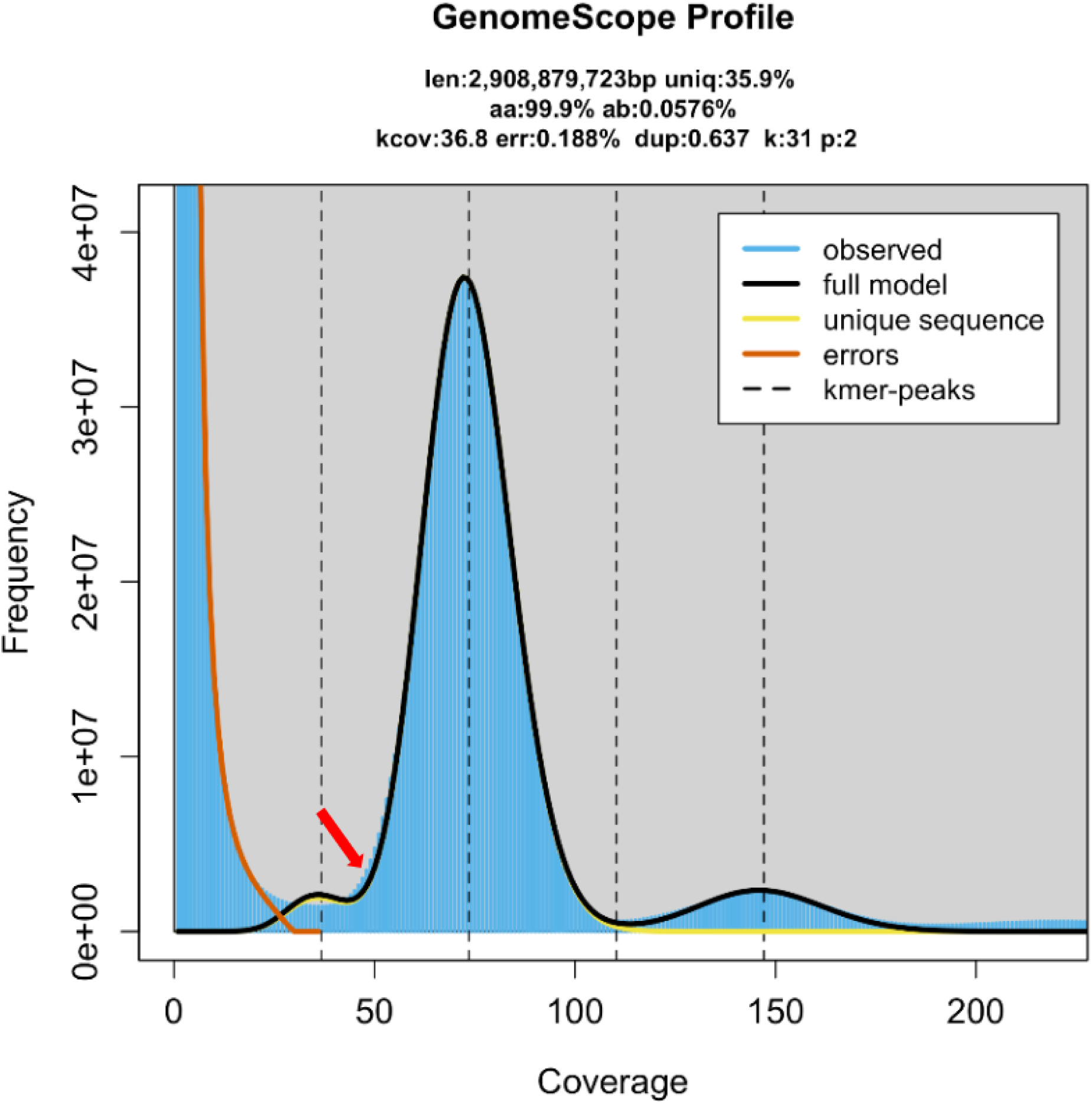
GenomeScope profile of *Alopecurus aequalis* HiFi reads using GenomeScope.FK. K-mer count distribution (in blue) from the HiFi reads showing a single homozygous peak at a coverage of 73, and a peak representing repeats at approximate coverage 150. The summary above the plot shows the estimated genome size in bp (len), unique fraction (uniq), homozygous fraction (aa), heterozygous fraction (ab), k-mer coverage of the heterozygous peak (kcov), PCR error estimation (err), PCR duplication estimation (dup), k-mer length (k), and ploidy (p). The red arrow indicates where the observed deviates from the model representing the heterozygous part of the genome.

### Genome assembly

Hifiasm^31^ v0.16.1 was used to generate a contig assembly and the homozygous coverage was specified as 73, corresponding to the position of the homozygous peak in the GenomeScope plot generated from the reads (Figure 1). Hifiasm generates assembly graphs for primary contigs and haplotype 1 and 2.

`hifiasm -o fx_run1.asm -t 64 --hom-cov 73 0.5_hifi_reads.fastq`

Reads assembled into 1,034 primary contigs with a total size of 2.9 Gb and a contig-N50 of 374.7 Mb (Table 2). The assembly size was similar to the k-mer based estimate and smaller than the flow-cytometry estimate. The total size of the individual haplotypes was similar to the primary contigs but the contiguity was lower (348 Mb and 244 Mb for haplotypes 1 and 2 respectively).

**Table 2:** Contiguity and completeness statistics from the contig assembly of *Alopecurus aequalis* showing primary contigs and the two assembled haplotypes. BUSCO abbreviations are C: Complete, S: Single copy, D: Duplicated, F: Fragmented, M: Missing, n: total BUSCO genes.

|  | Primary contigs | Haplotype 1 | Haplotype 2 |
| --- | --- | --- | --- |
| **Total bases (bp)** | 2,895,609,631 | 2,856,592,317 | 2,873,613,452 |
| **GC content (%)** | 45.57 | 45.55 | 45.58 |
| **Contig number** | 1,034 | 1,067 | 943 |
| **Longest contig (bp)** | 476,254,116 | 400,977,190 | 373,165,844 |
| **Contig N50 (bp)** | 374,747,030 | 347,700,133 | 244,747,959 |
| **Contig L50** | 4 | 4 | 5 |
| **Contig N90 (bp)** | 57,656,729 | 27,353,843 | 41,286,451 |
| **Contig L90** | 9 | 16 | 15 |
| **k-mer completeness (%)** | 98.0 | 97.7 | 97.9 |
| **BUSCO** | C:93.9%<br>[S:89.9%, D:4.0%],<br>F:1.6%, M:4.5%,<br>n:4896 | C:94.0%<br>[S:89.9%, D:4.1%],<br>F:1.5%, M:4.5%,<br>n:4896 | C:94.1%<br>[S:89.8%, D:4.3%],<br>F:1.5%, M:4.4%,<br>n:4896 |

The contiguity of this contig assembly was exceptionally high with the 13 longest contigs comprising 98% of the assembly. To determine whether these contigs represented whole chromosome arms we aligned contigs to the assembly of *A. myosuroides*^18^ and *H. vulgare*^32^ ‘Morex’ V3. Ten contigs were identified that represented chromosome or chromosome-arm level sequences. We also found two contigs with telomere sequences (TTTAGGG) on both ends and seven with telomere sequences on one end indicating that we had assembled chromosome or chromosome-arm level sequences. The low heterozygosity in *A. aequalis* estimated by GenomeScope was confirmed by identifying single-copy k-mers from a comparison of both haplotypes to the reads (Figure 3).

**Figure 3:**
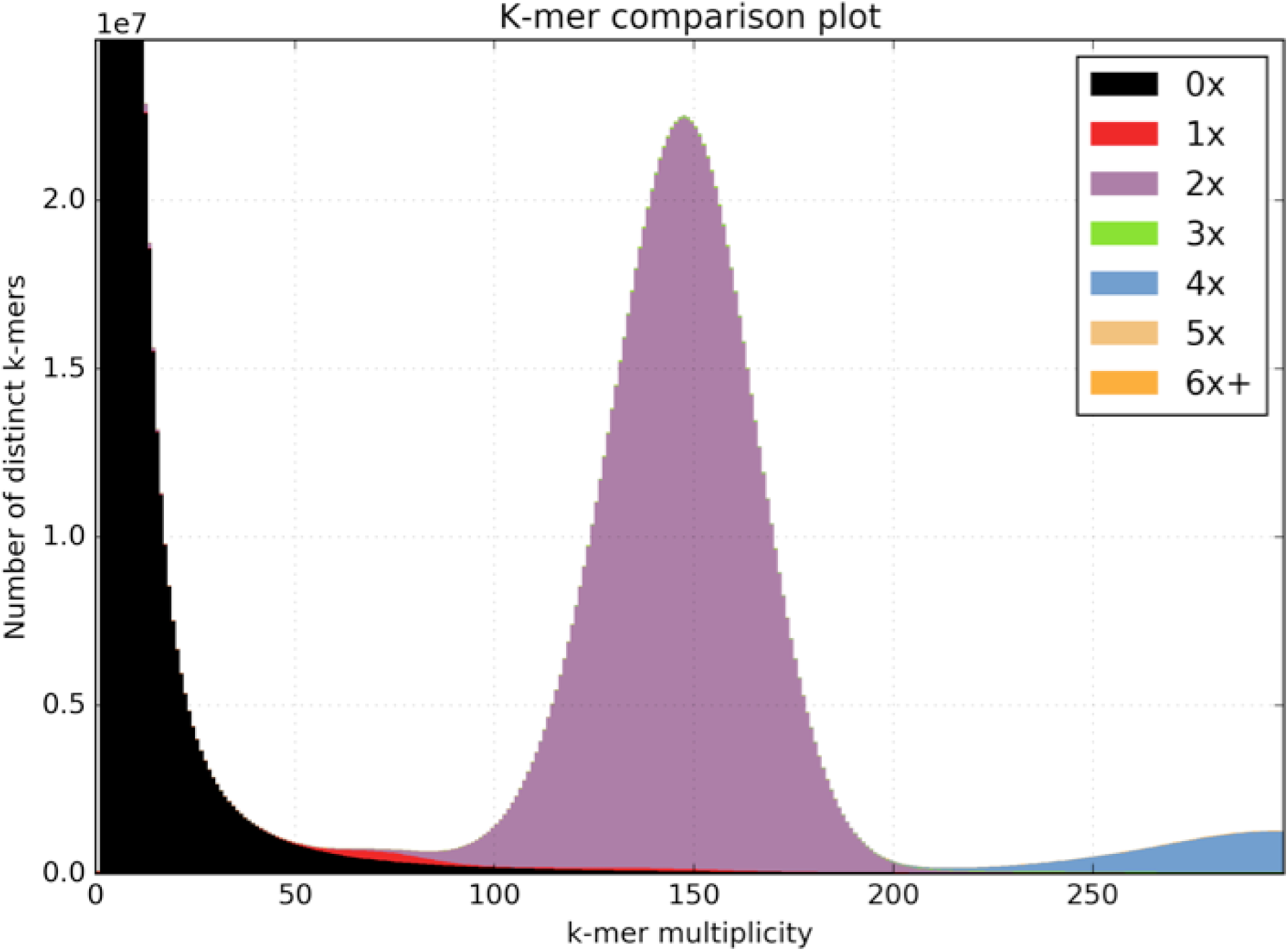
Content differences between the two assembled haplotypes of *Alopecurus aequalis.* K-mer copy-number spectra from KAT^33^ comparing both haplotypes to the reads. The majority of the kmers are present twice (once in each haplotype: purple peak), the red single-copy region represents kmers that exist in a single haplotype.

Contig scaffolding was performed using the Illumina paired-end Omni-C reads. The Arima Genomics Hi-C mapping pipeline^34^ was used to map the reads to the contigs before scaffolding with YaHS^35^. The mapping pipeline uses BWA^36^ (v.0.7.12) to map reads 1 and 2 separately before combining them into a single BAM file which is then deduplicated using Picardtools^37^ MarkDuplicates (v.2.25.7). The deduplicated BAM file was used as input to YaHS. This generated seven chromosome-level scaffolds, in agreement with previous cytogenetic studies^27^, and 421 sequences not incorporated into scaffolds.

To validate the scaffolding, Omni-C reads were mapped back to the scaffolds using BWA and the resulting contact map visualised using PretextView^38^. This identified one scaffold as retained haplotypic duplication and indicated a potential join between two larger scaffolds (scaffold_7 and scaffold_8). It also highlighted that many of the smaller scaffolds represented contamination. We removed the scaffold representing haplotypic duplication and joined scaffolds 7 and 8 into a larger scaffold. The final contact map is shown in Figure 4.

**Figure 4:**
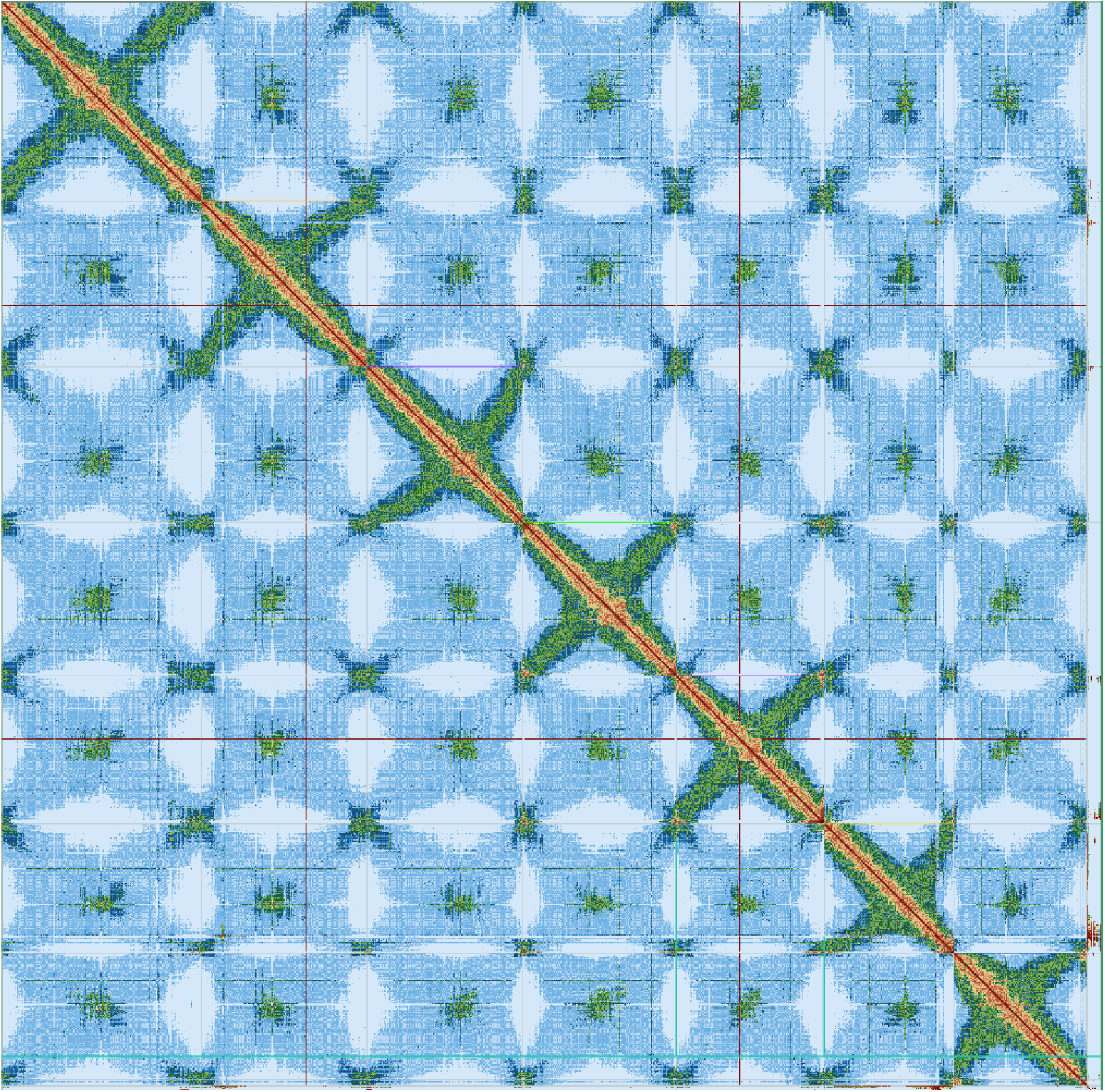
The PretextView contact map for *Alopecurus aequalis* after scaffolding and curation. Seven chromosomes in descending size order are represented by the diagonal red line with centromeric repeats showing as green regions. This map was generated after removal of one scaffold identified as a haplotypic duplication and the merging of scaffold_7 and scaffold_8.

Contaminated short sequences were identified and removed using NCBI BLAST+^39^ megablast (v2.12.0). The final assembly contained seven chromosome-level sequences and 145 shorter sequences not included in scaffolds. The chromosome-level scaffolds represented 2.8 Gb of sequence (99.5% of the assembly) and the total size of the 145 unassigned scaffolds was 15 Mb. Chromosome-level scaffolds were labeled in descending size order (Table 3).

**Table 3:** Final scaffold lengths and total assembly size for *Alopecurus aequalis*.

| Scaffold name | Length (bp) |
| --- | --- |
| chr_1 | 521,103,429 |
| chr_2 | 432,403,959 |
| chr_3 | 408,022,704 |
| chr_4 | 400,977,190 |
| chr_5 | 386,775,718 |
| chr_6 | 345,309,186 |
| chr_7 | 339,394,970 |
| Total | 2,833,987,156 |

### Genome annotation

Annotation was performed on contigs due to the high contiguity of the assembly at this stage. The annotation was lifted over to the scaffolds once they had been generated.

#### Repeat identification

Repeat annotation was performed using the EI-Repeat pipeline^40^ (v1.3.4) which uses third party tools for repeat calling. RepeatModeler^41^ (v1.0.11) was used for *de novo* identification of repetitive elements from the assembled *A. aequalis* genome using a customised repeat library. Unclassified repeats were searched in a custom BLAST database of organellar genomes (mitochondrial and plastid sequences from Pooideae in the NCBI nucleotide division) and any repeat families matching organellar DNA were also hard-masked. RepeatMasker^42^ (v4.0.72) with a RepBase^43^ embryophyte library and the customised RepeatModeler library was used to identify repeats. A total of 79% of the assembly was identified as repetitive and masked.

#### Reference guided transcriptome reconstruction

Gene models were derived from the RNA-Seq reads, Iso-Seq transcripts (HQ+LQ) and FLNC reads using the REAT transcriptome workflow^21^. HISAT2^44^ (v2.2.1) was used to align short reads and Iso-Seq transcripts aligned with minimap2^45^ (v2.18-r1015), maximum intron length was set as 50,000bp and minimum intron length to 20bp. Iso-Seq alignments were required to meet 95% coverage and 90% identity. High-confidence splice junctions were identified using Portcullis^46^ (v1.2.4). RNA-Seq Illumina reads were assembled for each tissue (root, shoot) with StringTie2^47^ (v2.1.5) and Scallop^48^ (v0.10.5), while FLNC reads were assembled using StringTie2. Gene models were derived from the RNA-Seq assemblies and Iso-Seq/FLNC alignments with Mikado^49^. Mikado was run with all Scallop, StringTie2, Iso-Seq and FLNC alignments and a second run with only Iso-Seq and FLNC alignments.

#### Cross-species protein alignment

Protein sequences from 11 Poaceae species (Table 4) were aligned to the *A. aequalis* assembly using the REAT Homology workflow^21^ which aligns proteins with spaln^50^ (v2.4.7) and miniprot^51^ (v0.3). The aligned proteins from both methods were clustered into loci and a consolidated set of gene models were derived via Mikado^49^.

**Table 4.** List of species used for cross species protein alignment. Evidence guided gene prediction.

| Organism Scientific Name | Assembly Accession |
| --- | --- |
| *Sorghum bicolor* | GCF_000003195.3 |
| *Brachypodium distachyon* | GCF_000005505.3 |
| *Setaria italica* | GCF_000263155.2 |
| *Oryza sativa* | GCF_001433935.1 |
| *Lolium rigidum* | GCF_022539505.1 |
| *Panicum hallii* | GCF_002211085.1 |
| *Lolium perenne* | GCF_019359855.1 |
| *Panicum virgatum* | GCF_016808335.1 |
| *Zea mays* | GCF_902167145.1 |
| *Hordeum vulgare* | GCF_904849725.1 |
| *Triticum aestivum* | iwgsc_refseqv2.1 (HC genes) |

Evidence guided annotation of protein coding genes was carried out using the REAT prediction workflow making use of the repeat annotation, RNA-Seq alignments, transcript assembly and alignment of protein sequences. The pipeline has four main steps:

1. The REAT transcriptome and homology Mikado models are categorised based on alignments to uniprot magnoliopsida proteins to identify models with likely full-length CDS and which meet basic structural checks. A subset of gene models is then selected from the classified models and used to train the AUGUSTUS^52^ gene predictor.
2. Augustus is run with extrinsic evidence generated in the REAT transcriptome and homology runs (repeats, protein alignments, RNA-Seq alignments, splice junctions, categorised Mikado models). Three evidence guided AUGUSTUS predictions are created using alternative bonus scores and priority based on evidence type.
3. AUGUSTUS models, REAT transcriptome / homology models, protein and transcriptome alignments are provided to EVidenceModeler^53^ (EVM) to generate consensus gene structures.
4. EVM models are processed through Mikado to add UTR features and splice variants.

#### Projection of gene models from *A. myosuroides*

*A. myosuroides* gene models^18^ were projected onto the *A. aequalis* assembly with Liftoff-1.5.1^54^, only those models that transferred fully with no loss of bases and identical exon/intron structure were retained (ei-liftover pipeline^55^).

#### Gene model consolidation

The final set of gene models was selected using Minos^56^, a pipeline that generates and utilises metrics derived from protein, transcript, and expression data sets to create a consolidated set of gene models. In this annotation, the following genes were filtered and consolidated into a single set of gene models:

1. The three alternative evidence guided Augustus gene builds derived from the REAT prediction run described earlier.
2. The EVM-Mikado gene models derived from the REAT prediction run described earlier.
3. The gene models derived from the REAT transcriptome runs described earlier.
4. The gene models derived from the REAT homology run described earlier.
5. The gene models derived from Liftoff.

Gene models were classified as biotypes protein_coding_gene, predicted_gene and transposable_element_gene, and assigned as high or low confidence based on the criteria below (results in Table 5):

a. **High confidence protein_coding_gene:** Any protein coding gene where any of its associated gene models have a BUSCO^57^ protein status of Complete/Duplicated OR have diamond^58^ (v0.9.36) coverage (average across query and target coverage) >= 90% against the listed protein datasets listed in Table 4 or Uniprot poaceae proteins. Alternatively they have average blastp coverage (across query and target coverage) >= 80% against the list protein datasets / uniprot poaceae AND have transcript alignment F1 score (average across nucleotide, exon and junction F1 scores based on RNA-Seq transcript assemblies) >= 60%.
b. **Low confidence protein_coding_gene**: Any protein coding gene where all of its associated transcript models do not meet the criteria to be considered as high confidence protein coding transcripts.
c. **High confidence transposable_element_gene**: Any protein coding gene where any of its associated gene models have coverage >= 40% against the combined interspersed repeats (see repeat identification section).
d. **Low confidence transposable_element_gene**: Any protein coding gene where all of its associated transcript models do not meet the criteria to be considered as high confidence and assigned as a transposable_element_gene.
e. **Low confidence predicted_gene**: Any protein coding gene where all of the associated transcript models do not meet the criteria to be considered as high confidence protein coding transcripts. In addition, where any of the associated gene models have average blastp coverage (across query and target coverage) < 30% against the protein datasets mentioned AND having a protein-coding potential score < 0.25 calculated using CPC2^59^ (v0.1).
f. **Low ncrna gene:** Any gene model with no CDS features AND a protein-coding potential score < 0.25 calculated using CPC2 0.1 and expression score > 0.6.
g. **Discarded models**: Any models having no BUSCO protein hit AND a protein alignment score (average across nucleotide, exon and junction F1 scores based on protein alignments) <0.2 AND a transcript alignment F1 score (average across nucleotide, exon and junction F1 scores based on RNA-Seq transcript assemblies) <0.2 AND a diamond coverage (target coverage) <0.3 AND Kallisto^60^ (v0.44) expression score <0.2 from across RNA-Seq reads OR having short CDS <30bps. Any ncRNA genes (no CDS features) not meeting the ncrna gene requirements (f) were also excluded.

**Table 5:** Minos classified gene models on the contig assembly of *Alopecurus aequalis*. Annotation liftover from contigs to scaffolds.

| **Biotype** | **Confidence** | **Genes** | **Transcripts** |
| --- | --- | --- | --- |
| protein_coding_gene | High | 35,149 | 53,355 |
| protein_coding_gene | Low | 20,304 | 21,383 |
| transposable_element_gene | Low | 3,617 | 3,691 |
| transposable_element_gene | High | 3,467 | 3,600 |
| predicted_gene | Low | 2,337 | 2,366 |
| ncrna_gene | Low | 575 | 1,313 |
| **Total** |  | **65,449** | **85,708** |

From the contig annotation (GFF3) we removed genes on contigs that were identified as contamination. Mikado was used to generate statistics from this GFF3 file. An AGP file was created to reflect the changes made during validation of the scaffolds and this was used as input to the *fromagp* function in the JCVI utility libraries^61^ (v0.8.12) to generate a chain file. Then the *gff* function of CrossMap^62^ (v0.3.4) was used to generate the liftover annotation (GFF3) from the modified contig GFF3. Mikado and GFFRead^63^ (v0.12.2) were used to check that all gene models had transferred correctly. A total of 33,758 high-confidence protein coding genes were annotated with a mean transcript length of 2,151 bp (Table 6). An additional 19,331 genes were classified as low-confidence.

**Table 6:** Genome annotation statistics for *Alopecurus aequalis*. High-confidence protein coding genes (HC), low-confidence protein coding genes (LC) and both sets combined (HC+LC).

|  | **HC** | **LC** | **HC+LC** |
| --- | --- | --- | --- |
| **Number of genes** | 33,758 | 19,331 | 53,089 |
| **Number of transcripts** | 51,946 | 20,409 | 72,355 |
| **Transcripts per gene** | 1.54 | 1.06 | 1.36 |
| **Number of monoexonic genes** | 7,933 | 6,035 | 13,968 |
| **Number of monoexonic transcripts** | 8,423 | 6,106 | 14,529 |
| **Transcript mean size cDNA (bp)** | 2,151.48 | 1,262.47 | 1,900.72 |
| **Transcript median size cDNA (bp)** | 1,906.00 | 1,103.00 | 1,653.00 |
| **Min cDNA length (bp)** | 156 | 159 | 156 |
| **Max cDNA length (bp)** | 17,668 | 8,838 | 17,668 |
| **Total exons** | 328,060 | 61,029 | 389,089 |
| Mean number of exons per transcript | 6.32 | 2.99 | 5.38 |
| Exon mean size (bp) | 340.67 | 422.19 | 353.46 |
| CDS mean size (bp) | 249.71 | 326.41 | 261.35 |
| Transcript mean size CDS (bp) | 1473.53 | 876.93 | 1,305.25 |
| Transcript median size CDS (bp) | 1,248 | 690.00 | 1,095.00 |
| Min CDS length (bp) | 156 | 138 | 138 |
| Max CDS length (bp) | 16,185 | 8,337 | 16,185 |
| Intron mean size (bp) | 448.30 | 968.72 | 515.04 |
| 5' UTR mean size (bp) | 262.00 | 141.15 | 227.91 |
| 3' UTR mean size (bp) | 415.94 | 224.39 | 367.55 |

#### Functional annotation

All the proteins were annotated using AHRD^64^ (v.3.3.3). Sequences were BLASTed against the reference proteins (Arabidopsis thaliana TAIR10, TAIR10_pep_20101214_updated.fasta.gz - https://www.araport.org) and the UniProt viridiplantae sequences (data download 06-May-2023), both Swiss-Prot and TrEMBL datasets^65^. Proteins were BLASTed (v2.6.0; blastp) with an value of 1e-5. We also provided InterProScan^66^ (v5.22.61) results to AHRD. We adapted the standard AHRD configuration file (test/resources/ahrd_example_input_go_prediction.yml), distributed with AHRD, changing the following:

1. We included the GOA mapping from uniprot (ftp://ftp.ebi.ac.uk/pub/databases/GO/goa/UNIPROT/goa_uniprot_all.gaf.gz) as the parameter ’gene_ontology_result’,
2. We included the interpro database (ftp://ftp.ebi.ac.uk/pub/databases/interpro/61.0/interpro.xml.gz) and provided as parameter ’interpro_database’,
3. We changed the parameter ’prefer_reference_with_go_annos’ to ’false’
4. The blast database specific weights used were:

- swissprot: weight=100, description_score_bit_score_weight=0.2
- trembl: weight=50, description_score_bit_score_weight=0.4
- tair: weight=50, description_score_bit_score_weight=0.4

### Genome overview

The *A. aequalis* genome contains seven chromosomes ranging in size from 521 to 339 Mb (Figure 5). Gene density is highest at the distal ends of the chromosomes with very few genes in the centromeric regions. The distribution of transposable elements is quite even over the length of the chromosomes. The distribution of Ty3/*Gypsy* LTR retrotransposons (LTR RTs) is highest in gene-poor regions with fewer found in the gene-rich distal ends of the chromosomes. Ty1/*Copia* LTR RTs are distributed throughout the chromosomes with fewer in centromeric regions.

**Figure 5:**
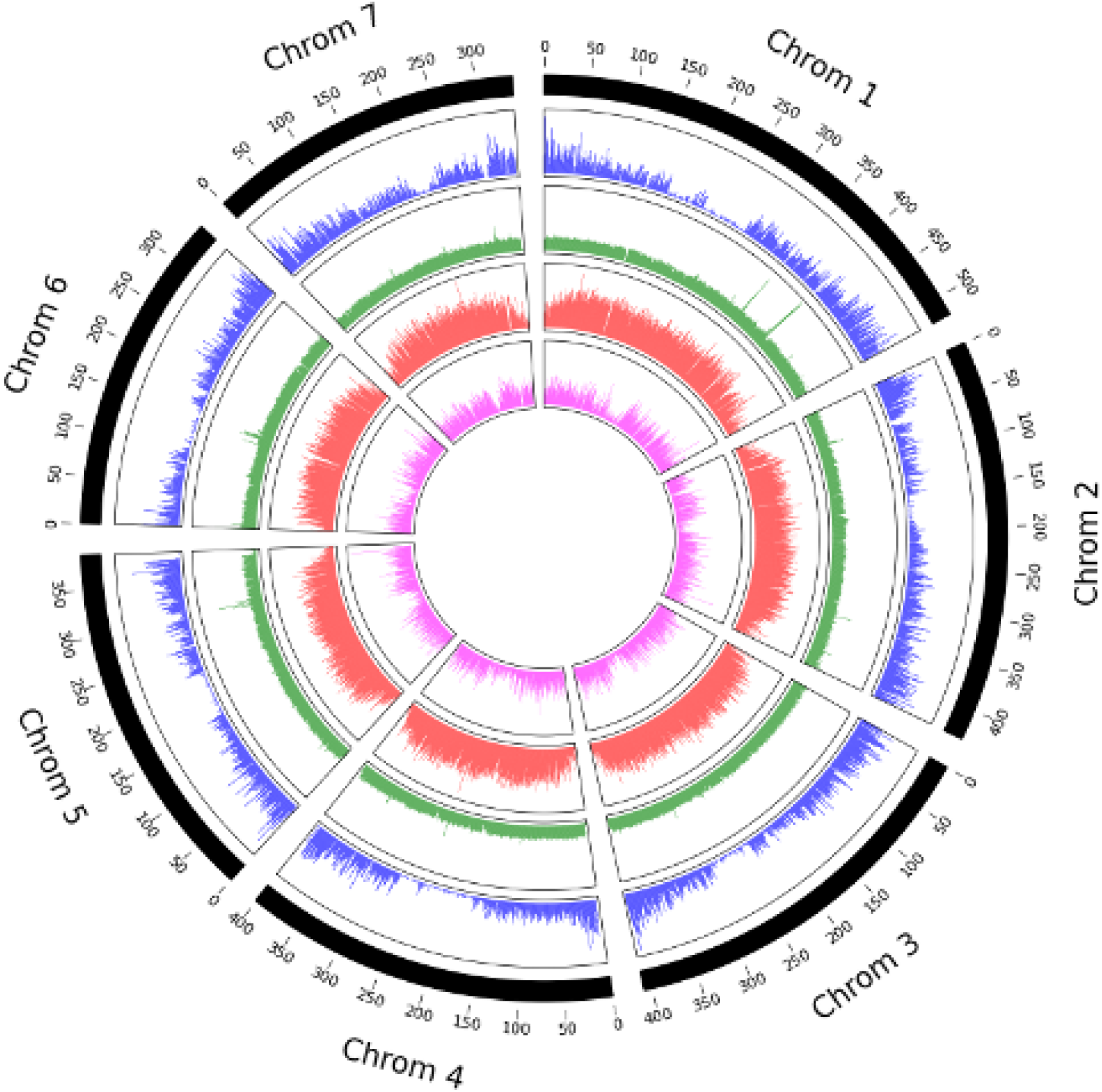
Overview of the *Alopecurus aequalis* genome. Distribution of high-confidence protein-coding genes (blue), distribution of transposable elements (green), distribution of Ty3/*Gypsy* long-terminal repeats retrotransposons (LTR-RTs) (red) and distribution of Ty1/*Copia* LTR RTs (pink).

### Comparative genomics analysis

GENESPACE^67^ was used to assess synteny between *A. aequalis, A. myosuroides*^18^ and *Hordeum vulgare* cultivar *‘*Morex’^68^ to identify large structural rearrangements. Although a second *A. myosuroides* genome is available^19^, our analyses showed no differences between the two versions. Therefore, we used the genome from Cai et al.^18^ in all of our analyses.

The seven chromosomes of *A. aequalis* are more similar in structure to *H. vulgare* (Figure 6A). Six of the seven chromosomes show a high level of synteny to *H. vulgare* with *A. aequalis* chromosome 1 comprising two syntenic blocks from *H. vulgare* chromosomes 4H and 5H. There is also evidence of a small translocation between *A. aequalis* chromosome 6 to *H. vulgare* chromosomes 4H.

**Figure 6:**
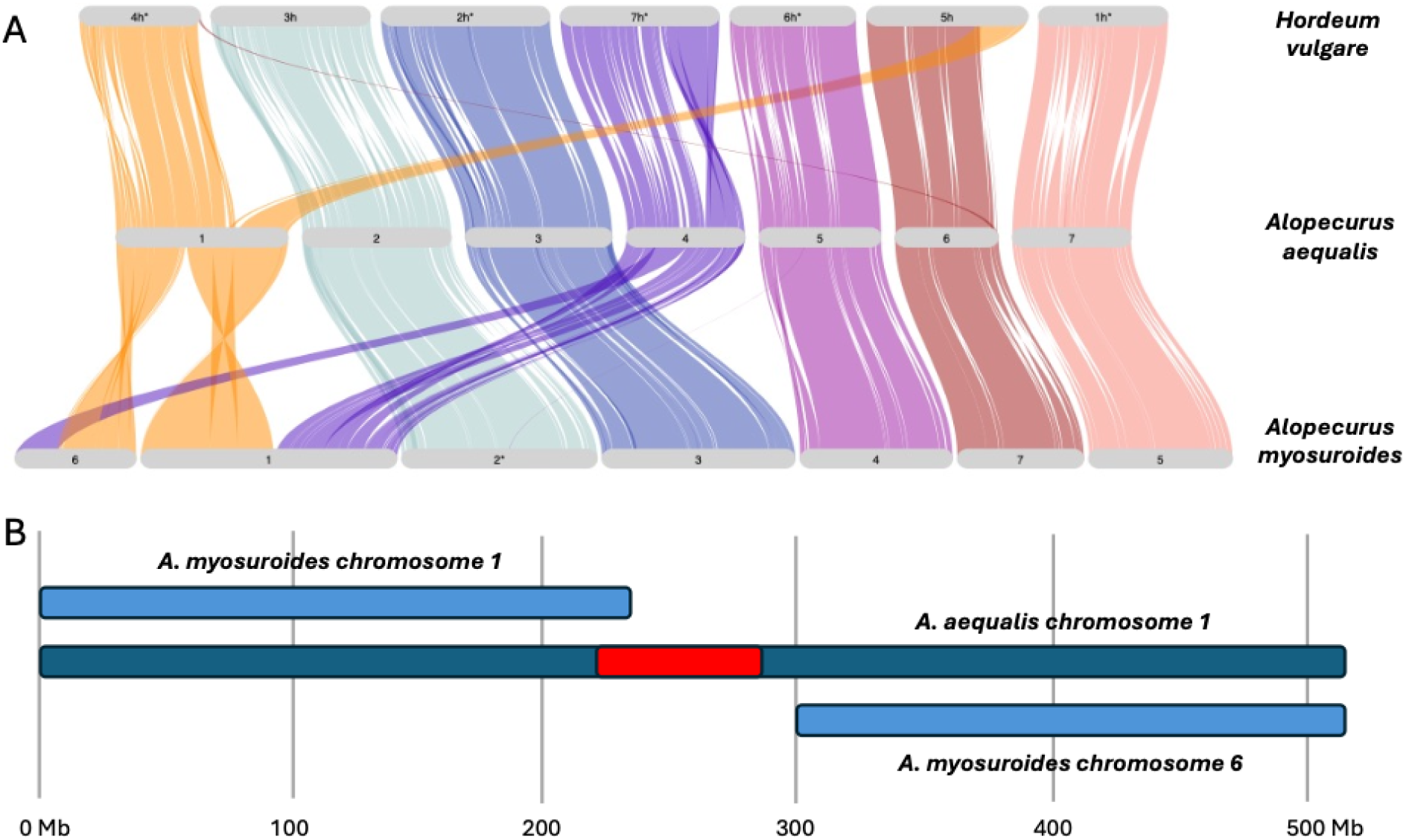
Whole genome comparison between *Hordeum vulgare*, *Alopecurus aequalis* and *Alopecurus myosuroides*. A: GENESPACE comparison showing synteny between the 3 species (* indicates that the chromosome has been reversed). B: Detailed alignment showing the position of the *A. aequalis* chromosome 1 centromere (in red) in relation to the breakpoint.

Five chromosomes of *A. aequalis* show a high level of synteny to *A. myosuroides.* The chromosome arms of *A. aequalis* chromosome 1 are syntenic to two regions of *A. myosuroides* chromosomes 1 and 6. The break in synteny appears to occur in the centromeric region of *A. aequalis* chromosome 1, estimated to be between 220 and 290 Mb according to the Hi-C contact map (Figure 4) which also corresponds to the drop in gene density in this region of the chromosome (Figure 5). More detailed alignments show *A. aequalis* chromosome 1 aligns to *A. myosuroides* chromosome 1 from 0-240 Mb and to *A. myosuroides* chromosome 6 from 300-521 Mb (Figure 6B).

Chromosome 4 of *A. aequalis* is syntenic to two large regions on *A. myosuroides* chromosomes 1 and 6. This relationship is more complex, showing several internal rearrangements within the larger syntenic region. It should be noted that the differences between *A. aequalis* chromosomes 1 and 4 compared to *A. myosuroides* chromosomes 1 and 6 were evident in the contig stage of assembly.

### Genes involved in herbicide resistance

Cytochrome P450 genes have been identified as the main genes involved in non-target site herbicide resistance^6,7^. In *A. myosuroides*^18^, where the P450 gene family has significantly expanded compared to Arabidopsis and rice, 506 loci were identified, mainly on chromosomes 2 and 3. The functional annotation of *A. aequalis* identified 513 P450 genes, suggesting this species has also experienced an expansion of genes involved in non-target site herbicide resistance (Figure 7).

**Figure 7:**
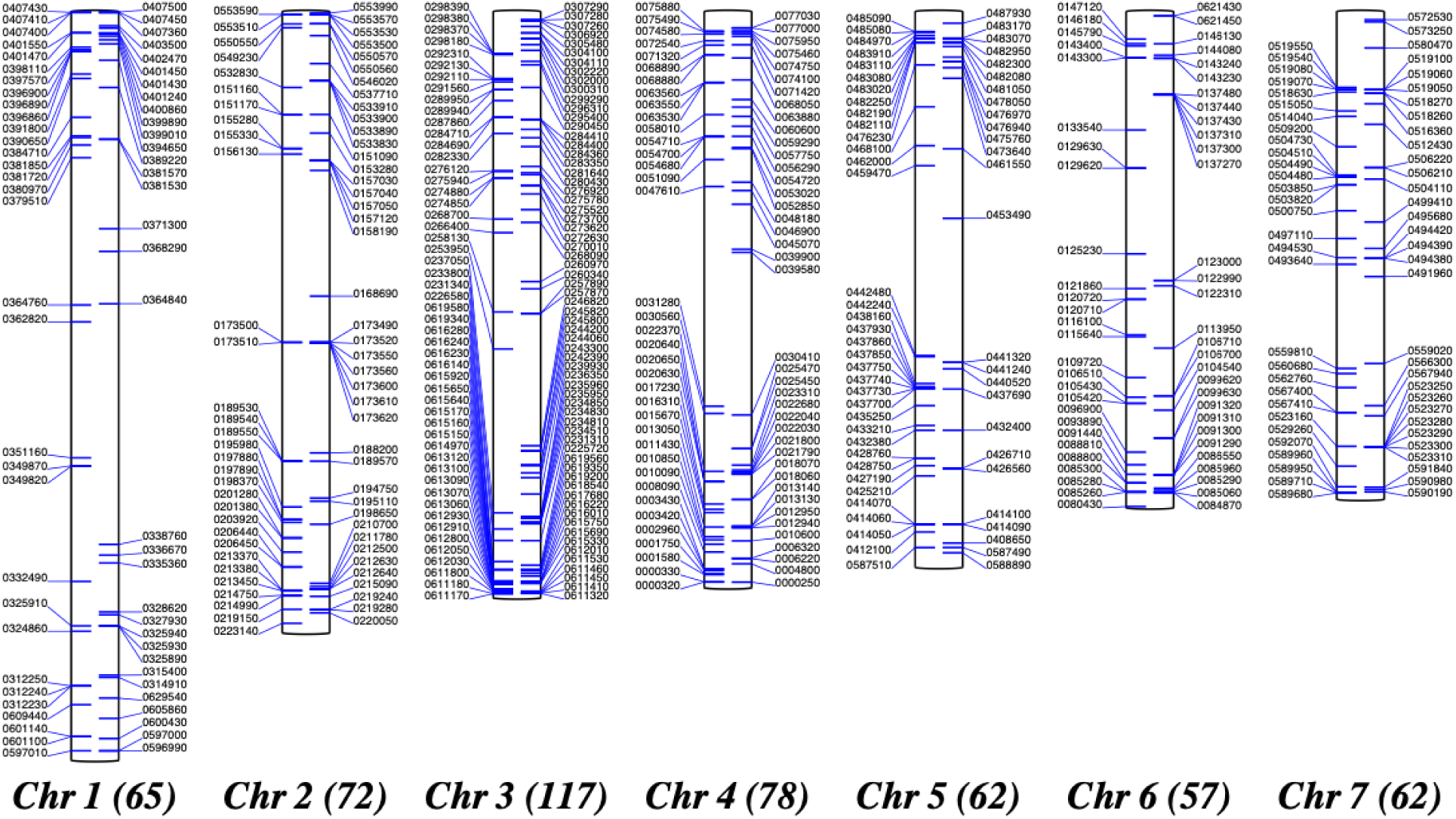
Locations of cytochrome P450 gene loci in *Alopecurus aequalis*. The number in brackets shows the total number of loci on that chromosome. Gene identifiers are shown without the Alaeq_EIv0.2_ prefix for clarity.

## Data Records

All read datasets used in the assembly and annotation of *Alopecurus aequalis* are available at the National Center for Biotechnology Information (NCBI) under Project ID PRJEB75647.

Genome assembly and annotation files are available from https://opendata.earlham.ac.uk/opendata/data/wright_sep2024_orange_foxtail/.

## Technical Validation

### Evaluating the quality of the genome assembly

BUSCO^57^ (v5.3.2) and Merqury^69^ (v1.3) were used to assess assembly completeness. BUSCO analysis showed 4,595 (93.9%) complete BUSCO genes from the poales_odb10 lineage dataset (total: 4,896) were found in the primary contigs. Of these, 4,401 were found as single-copy genes and 194 as duplicated genes. 77 BUSCO genes were fragmented and 224 were missing from the assembly. Merqury computed a k-mer completeness metric of 98.0% meaning that 98% of kmers from the reads are found in the assembly. The QV quality score was 61.6 corresponding to a base level accuracy of 99.9999%.

The Merqury spectra copy-number plot shows the majority of k-mers from the reads are found in the primary contigs only once at the expected coverage (the red region), and the majority of the low-coverage k-mers (originating from errors in the reads) are not present in the primary contigs (Figure 8).

**Figure 8:**
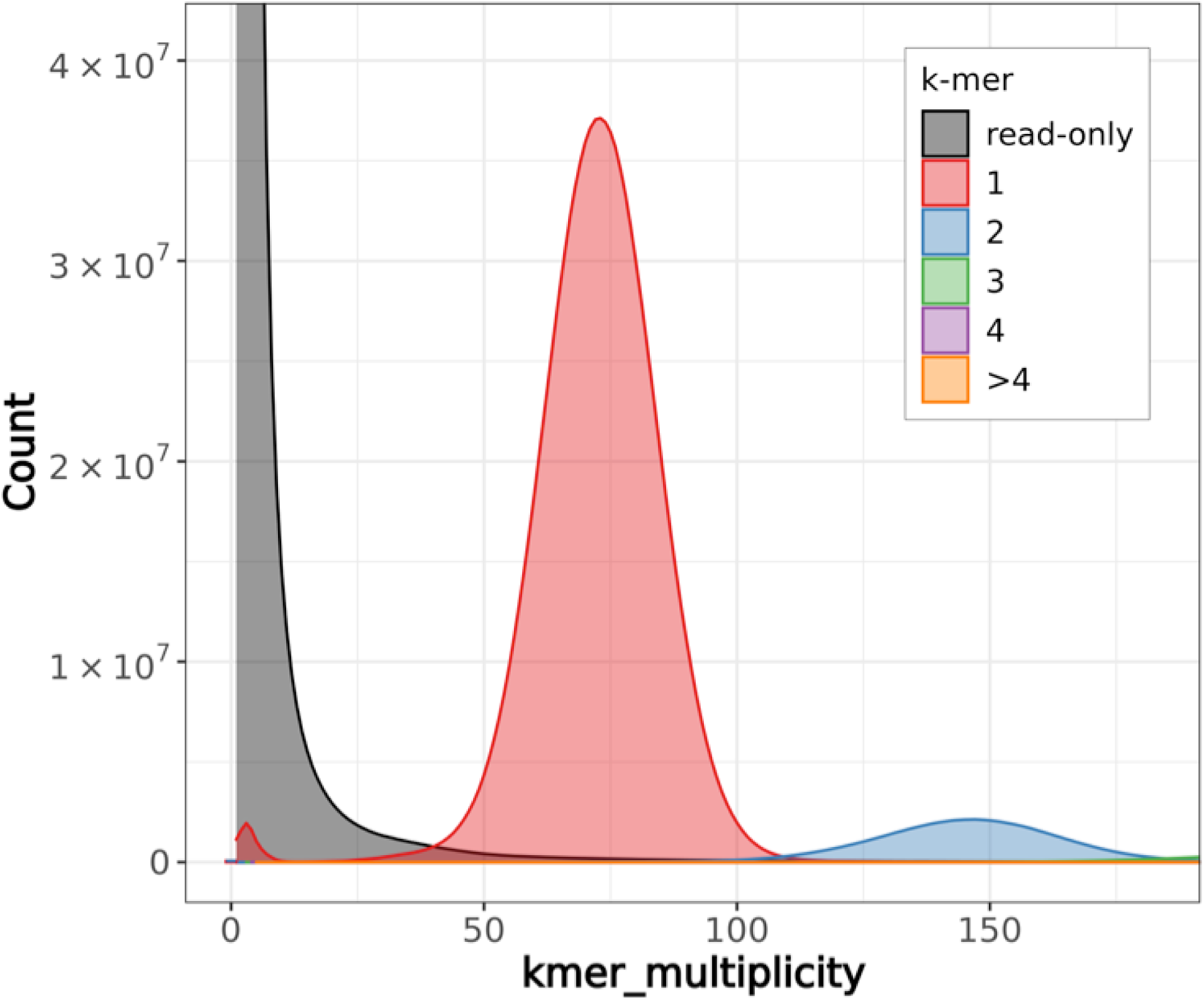
Spectra copy-number plot comparing k-mers from reads to k-mers in the primary contig assembly of *Alopecurus aequalis*.

After scaffolding primary contigs using Omni-C data, the Omni-C reads were mapped back to the scaffolds to identify erroneous joins between contigs. After removing a duplicated scaffold and joining two others we were satisfied that the assembly was of high-quality.

### Evaluating the quality of the genome annotation

BUSCO^57^ with the poales_odb10 lineage dataset was used to measure the completeness of the high-confidence protein-coding gene set. In total, 99.1% of BUSCO groups were marked as complete (4,855 out of 4,896), 92.1% were complete and single-copy. There were two fragmented and 39 missing BUSCO groups indicating that the high-confidence protein coding gene set represents the *A. aequalis* gene complement accurately. OMArk^70^ was used to assess the completeness and consistency of the *A. aequalis* annotation against gene families in the Pooideae clade (Figure 9). The high-confidence gene set was compared to Hierarchical Orthologous Groups (HOGs) to give an estimate of completeness which showed 93.7% of HOGs were found (83.0% single and 10.7% duplicated) with 6.4% missing. When the combined high and low confidence genes are compared to HOGs we find 95.6% of HOGs with 4.5% missing indicating that some low confidence genes represent valid HOGs. For the high confidence gene set, 93.8% of genes were consistent with gene families in the Pooideae clade, with 3.4% found in different lineages and 2.8% of unknown origin.

**Figure 9:**
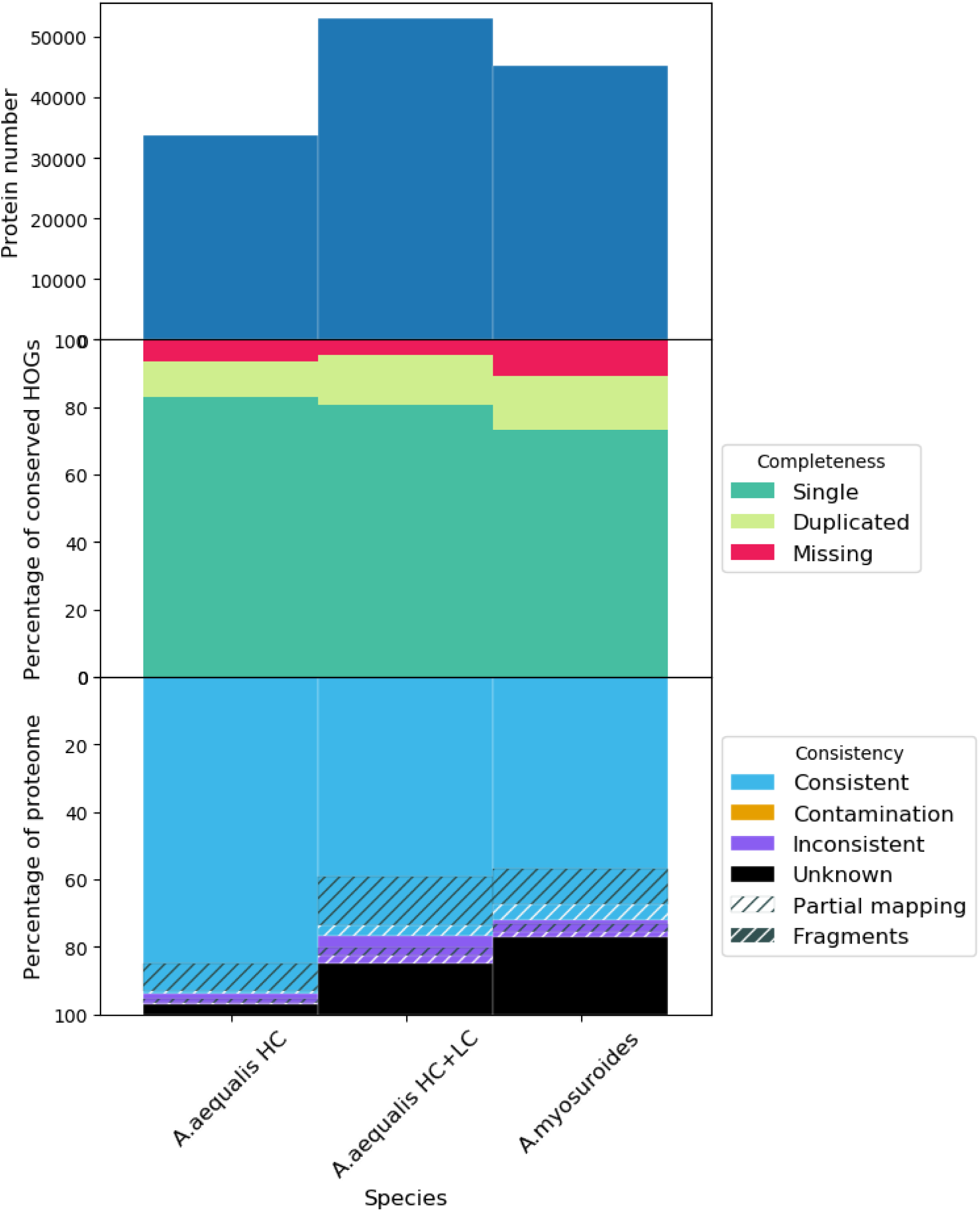
OMArk plot comparing protein number, completeness and consistency of the *Alopecurus aequalis* genome annotation (HC: high confidence protein-coding genes, LC: low confidence protein-coding genes). OMArk output from the *A. myosuroides* genome annotation is included for comparison.

We identified 11,505 fewer genes in *A. aequalis* compared to that reported for *A. myosuroides*^18^ (45,263), a difference likely caused by the different annotation methods used for each genome and the classification used in this pipeline to separate genes into high and low-confidence.

Running OMArk on the *A. myosuroides* annotation (Figure 9) shows more missing genes (10.8%) indicating a less complete annotation as well as more genes classified as “Unknown” (23.0%) indicating that many of the gene models included in the *A. myosuroides* annotation probably represent low confidence genes. The recent annotation of the closely related barley cultivar ‘Morex’ identified 32,787 high-confidence gene models^68^ which is more similar to the annotation presented here.

We also used OrthoFinder^71^ (v2.0.9) to cluster *A. aequalis* high-confidence genes and *A. myosuroides* genes into orthogroups. A total of 50,904 orthogroups were generated, 3,664 containing multiple genes, and 17,375 single-copy orthogroups. 9,660 orthogroups contained a single *A. aequalis* gene and 20,804 orthogroups contained a single *A. myosuroides* gene, also indicating that the *A. myosuroides* gene set contains many genes that are not present in the *A. aequalis* high-confidence gene set.

## Code Availability

All software used in this study was run according to instructions. The version and parameters are described in the methods. Anything not described in Methods was run with default parameters.

## Acknowledgements

The authors would like to thank the ERGA executive team, Ann Mc Cartney, Alice Mouton, and Guilio Formenti for initiating the ERGA project and for ongoing support. They also would like to acknowledge Faye Oddy, Richard Hull, Laura Crook, and members of the Rothamsted Research Horticulture and Controlled Environment teams for their support with bulking seed and maintaining plants. The authors acknowledge the support of the Biotechnology and Biological Sciences Research Council (BBSRC), part of UK Research and Innovation; Earlham Institute (EI) Strategic Programme Grant Decoding Biodiversity BBX011089/1 and its constituent work package BBS/E/ER/230002B (Decode WP2 Genome Enabled Analysis of Diversity to Identify Gene Function, Biosynthetic Pathways, and Variation in Agri/Aquacultural Traits), as well as the Core Capability Grant BB/CCG2220/1 and the work delivered via the Research Computing Groups who manage and deliver High Performance Computing at EI. Part of this work was delivered via the BBSRC-funded National Capability in Genomics and Single Cell Analysis (BBS/E/T/000PR9816) and National Bioscience Research Infrastructure in Transformative Genomics (BBS/E/ER/23NB0006) at Earlham Institute by members of the Technical Genomics and the Core Bioinformatics Groups. Rothamsted Research receives strategic funding from the Biotechnology and Biological Sciences Research Council of the United Kingdom (BBSRC) and the authors acknowledge support from the Smart Crop Protection Industrial Strategy Challenge Fund (grant no. BBS/OS/CP/000001) and the Growing Health Institute Strategic Programme (BB/X010953/1; BBS/E/RH/230003A). Part of this work was supported by Wellcome through the Darwin Tree of Life Discretionary Award (218328).

## Author contributions

D.M. and A.H. conceived the project and designed the study. C.H. and D.M. grew the plants and sampled them. R.P. performed the flow cytometry analysis and I.J.L. coordinated this analysis. H.R performed initial sample coordination and DNA extraction. F.F. developed methods for and prepared Dovetail Omni-C and RNA-seq libraries. N.I. developed methods for and prepared HiFi and Iso-Seq libraries. A.D. developed the method for and performed the DNA extraction for HiFi library preparation. S.H. prepared RNA-seq libraries. T.B. sequenced Illumina RNA-seq libraries, Dovetail Omni-C library, PacBio Iso-Seq libraries, and PacBio HiFi libraries. J.W. performed genome assembly, genome annotation liftover and comparative analyses. G.K. and D.S. performed genome annotation. J.Wo. assisted with the interpretation of the Hi-C contact map. C.W., K.G., and L.C. planned and supervised data production. S.L. and K.B. coordinated the project from sample submission to data delivery. S.M. managed the project with respect to ERGA. J.W., D.M., C.W. and D.S. wrote the initial manuscript draft and A.H., H.R., J.L., P.N. and I.J.L. contributed editing and improvements. All authors read and approved the final manuscript.

## Competing Interests

The authors declare no competing interests.

## Notes

### Competing Interest Statement

The authors have declared no competing interest.

